# One-to-one innervation of vocal muscles allows precise control of birdsong

**DOI:** 10.1101/2020.01.10.901561

**Authors:** I. Adam, A. Maxwell, H. Rössler, E.B. Hansen, M. Vellema, C.P.H. Elemans

## Abstract

The motor control resolution of any animal behavior is limited to the minimal force step available when activating muscles, which is set by the number and size distribution of motor units (MUs) and muscle specific force [1, 2]. Birdsong is an excellent model system for understanding sequence learning of complex fine motor skills [3], but we know surprisingly little how the motor pool controlling the syrinx is organized [4] and how MU recruitment drives changes in vocal output [5]. Here we combine measurements of syringeal muscle innervation ratios with muscle stress and an *in vitro* syrinx preparation to estimate MU size distribution and the control resolution of fundamental frequency (*f*_o_), a key vocal parameter, in zebra finches. We show that syringeal muscles have extremely small MUs, with 50% innervating ≤ 3, and 13 – 17% innervating a single muscle fiber. Combined with the lowest specific stress (5 mN/mm^2^) known to skeletal vertebrate muscle, small force steps by the major *f*_o_ controlling muscle provide control of 50 mHz to 4.2 Hz steps per MU. We show that the song system has the highest motor control resolution possible in the vertebrate nervous system and suggest this evolved due to strong selection on fine gradation of vocal output. Furthermore, we propose that high-resolution motor control was a key feature contributing to the radiation of songbirds that allowed diversification of song and speciation by vocal space expansion.

## Results & Discussion

Vocal communication is of paramount importance for reproduction and survival of songbirds and even drives speciation. This is clearly exemplified by sympatric species that are morphologically indistinguishable, but nevertheless fully separated solely by song [6, 7]. The potential to acoustically separate depends on the number of distinct sounds – or vocal space – that can be produced and perceived. The vocal space is set both by the range available to vary an acoustic feature, and in what steps, or resolution, the feature can be controlled within this range. Thus, both resolution and range expand the vocal space and may form a rich substrate for species diversification [8, 9]. While the range of an acoustic feature, for example *f*_o_, is typically limited by intrinsic constraints of the vocal organ [10-12], the ability to generate distinct *f*_o_ changes within these constraints is set by the resolution of the neural control. In contrast to the well-described neural circuitry underlying song learning [3], we know surprisingly little how the motor pool controlling the songbird syrinx is organized [4]. Furthermore, even though birdsong is often called a fine-motor skill [13, 14] and perturbed auditory feedback can drive small *f*_o_ changes [15, 16], the control resolution of acoustic features remains unknown.

Motor control of any behavior is limited to the minimal force step available when activating muscles, which is set by the number and size distribution of MUs and muscle fiber specific force generation [1]. A MU is the basic functional unit of skeletal muscle and consists of a motor neuron and the number of muscle fibers it innervates, aka the innervation number (*IN*). Variation in *IN*, muscle specific force and spike rate are the most significant factors to contribute to differences in MU force in skeletal muscles [1]. Currently, we do not know the minimal force step available in vocal motor control and how MU recruitment causes changes in vocal output in songbirds.

### The songbird syrinx motor pool has the highest control resolution possible

We quantified the mean *IN* (*IN*_mean_) of syringeal muscles in male zebra finches by counting *i)* the number of muscle fibers and *ii)* the number of axons in the supplying ipsilateral nerve (Fig 1A, B, Table S1, See Methods). The average total number of muscle fibers was 6995 ± 789 (n = 4), corroborating earlier reported ∼6730 [17]. The left side had significantly less muscle fibers (3234 ± 232, range: 2992 – 3503, n = 4) than right (3794 ± 334, range 3586 – 4286, n = 4) (Welch t-test: t = −2.8, df = 5.4, p-value = 0.04). The number of axons in the tracheosyringeal branch of the hypoglossal nerve (NXIIts), assumed to represent the number of MUs (See Methods), was not significantly different between left (820 ± 187, range: 602 – 1162, n = 8) and right (790 ± 148, range: 677 – 1045, n = 5) (Welch t-test: t = 0.32, df = 10.2, p-value = 0.76) and lower but within range of the 1026 ± 126 (n = 6) reported earlier [18]. The individual ipsilateral *IN*_mean_ was not significantly different between left and right (Welch t-test: t = −1.5, df = 4.98, p = 0.18) and ranged from 3.85 to 5.67 (4.78 ± 0.75, n = 7). Vertebrate muscles are typically innervated by a few hundred MUs, with *INs* of several hundred muscle fibers [1]. The *IN*_mean_ of syringeal muscles is thus extremely low, and comparable only to, but still lower than, laryngeal [19] and extraocular [20] muscles.

**Figure 1.**
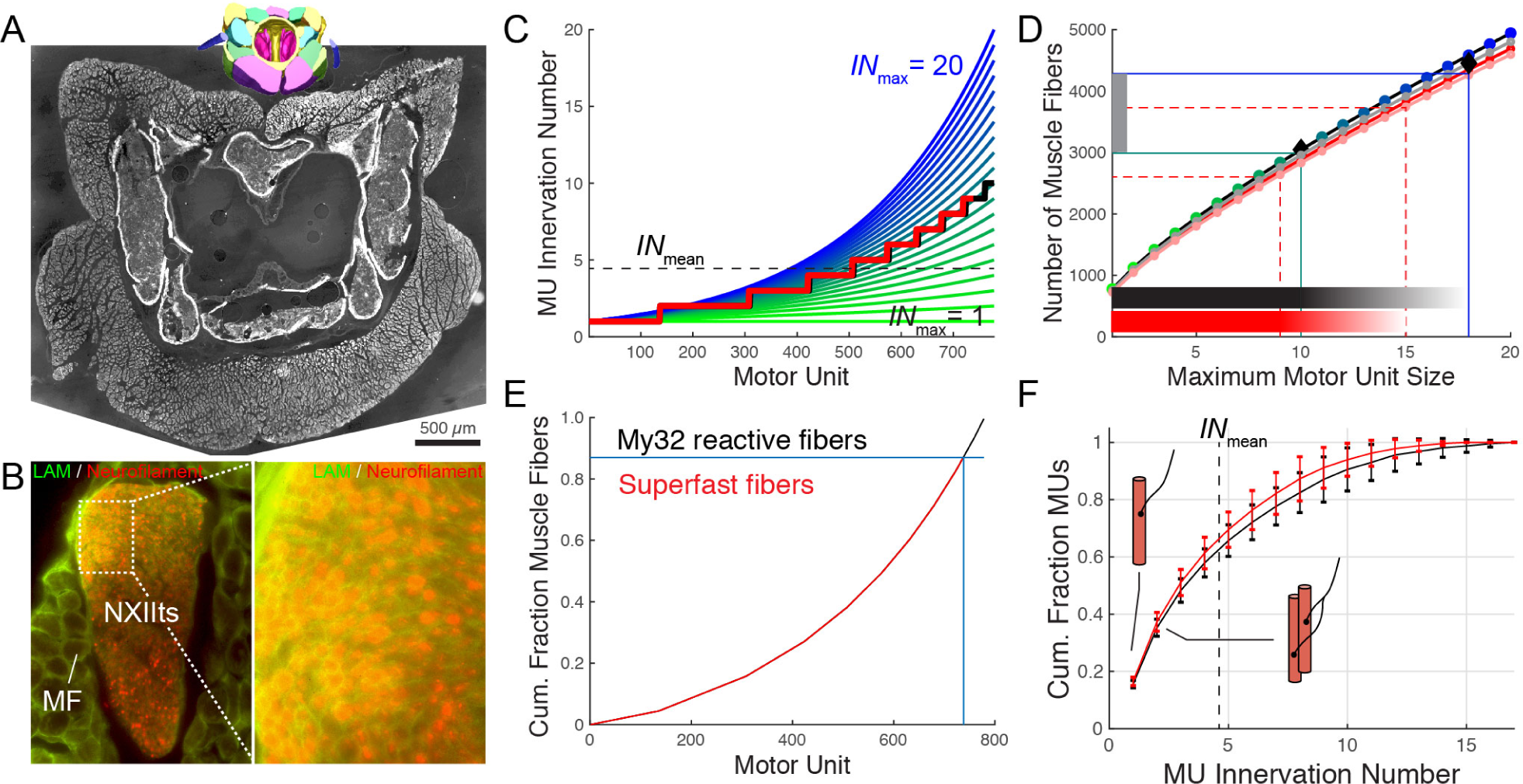
The songbird syrinx motor pool has the highest control resolution possible. A) Cross-sections through A) syrinx at mid-level (∼800 stitched images) and B) the tracheosyringeal branch of the hypoglossal nerve (NXIIts). C) *IN* distribution as function of MU with *IN*_max_ ranging from 1 (green) to 20 (blue). Black lines indicate the average *IN*_mean_ (horizontal dotted line) and distributed *IN* (stepped line) for the individual with the smallest *IN*_mean_ (3.84). The red line (also next panels) indicates the distribution for only superfast muscle fibers. D) The number of muscle fibers (horizontal lines) provides *IN*_max_ per individual. Colors correspond to *IN*_max_ lines in panel C. Shown are the individuals with smallest (3.84, green) and largest *IN*_mean_ (5.67, blue). Shaded horizontal bars indicate *IN* range. E) Cumulative fraction of muscle fibers with MU number. Of all muscle fibers 87% do not react to my32 and are most likely of the superfast fiber type (red). F) Cumulative fraction of MU as a function of *IN* for all (black) and superfast fibers (red). Data is presented as mean ± SD.

However, critical to muscle function is not the *IN*_mean_ among MUs, but *IN* distribution within a given motor pool [1]. *IN* distribution sets the smallest force step possible and the motor pool proportions that innervate different muscle fiber types [1]. We estimated the frequency distribution of MU size within syringeal muscles by exploiting the finding that MU size distribution and MU force are consistently skewed for all muscles studied to date [21-23] (See Methods): the majority of MUs are small and only few are large. Using our empirical data on the number of MUs and muscle fibers, we systematically varied (Fig 1C) the size of the largest MU to find the value where the predicted number matches the counted number of muscle fibers (Fig 1D). We found that syringeal MUs are extremely small and range in size from 1 to 13.2 ± 2.8 (range of maximal MU size: 10 – 15, n = 7) muscle fibers. As a consequence, half of all syringeal MUs (49 ± 4%, n = 7) innervate as few as ≤ 3 muscle fibers, and 13 ± 3% (n = 7) of the MUs innervate a single muscle fiber (Fig 1G). About 87% of all muscle fibers in adult male zebra finches seem of the superfast phenotype [17]. Including only superfast fibers (red lines in Fig1 E-F, see Methods), shifts the MU size distribution curve slightly to the left. Superfast syringeal MUs contain 1 to 11.6 ± 2.6 muscle fibers (range of maximal MU sizes 10 – 15, n = 7), with 52 ± 5% (n = 7) innervating as few as ≤ 3 muscle fibers and almost a fifth (17 ± 2%, (n = 7)) of all motor neurons innervating a single superfast syringeal muscle fiber. Such one-to-one innervation has to our knowledge only been reported for monkey extraocular muscle [24] and one motor neuron of the mouse interscutularis muscle [25] and provides the nXIIts motor pool with the highest control resolution possible in the vertebrate nervous system.

### Songbird syringeal muscles generate the lowest stress

The other determinant of MU force is set by muscle-specific intrinsic properties, particularly the stress (*P*), a fiber generates. Stress ranges from the peak stress during the all-or-none twitch contraction (*P*_tw_) caused by a single motor neuron spike, up to the maximum stress during tetanic contraction (*P*_o_) caused by spike trains of the motor neuron. Because syringeal muscles are superfast muscles that can power movement up to 200 – 250 Hz [26], they trade force for speed as a fundamental architectural trait [27, 28] and are expected to have low *P*_o_. Indeed, pigeon syringeal muscles generate a *P*_o_ as low as 18 – 50 mN/mm^2^ [29], 5 – 10 times lower as regular skeletal muscle fibers (∼150 – 300 mN/mm^2^) [30, 31] (Fig 2), but the *P*_o_ of songbird syringeal muscles is unknown.

**Figure 2.**
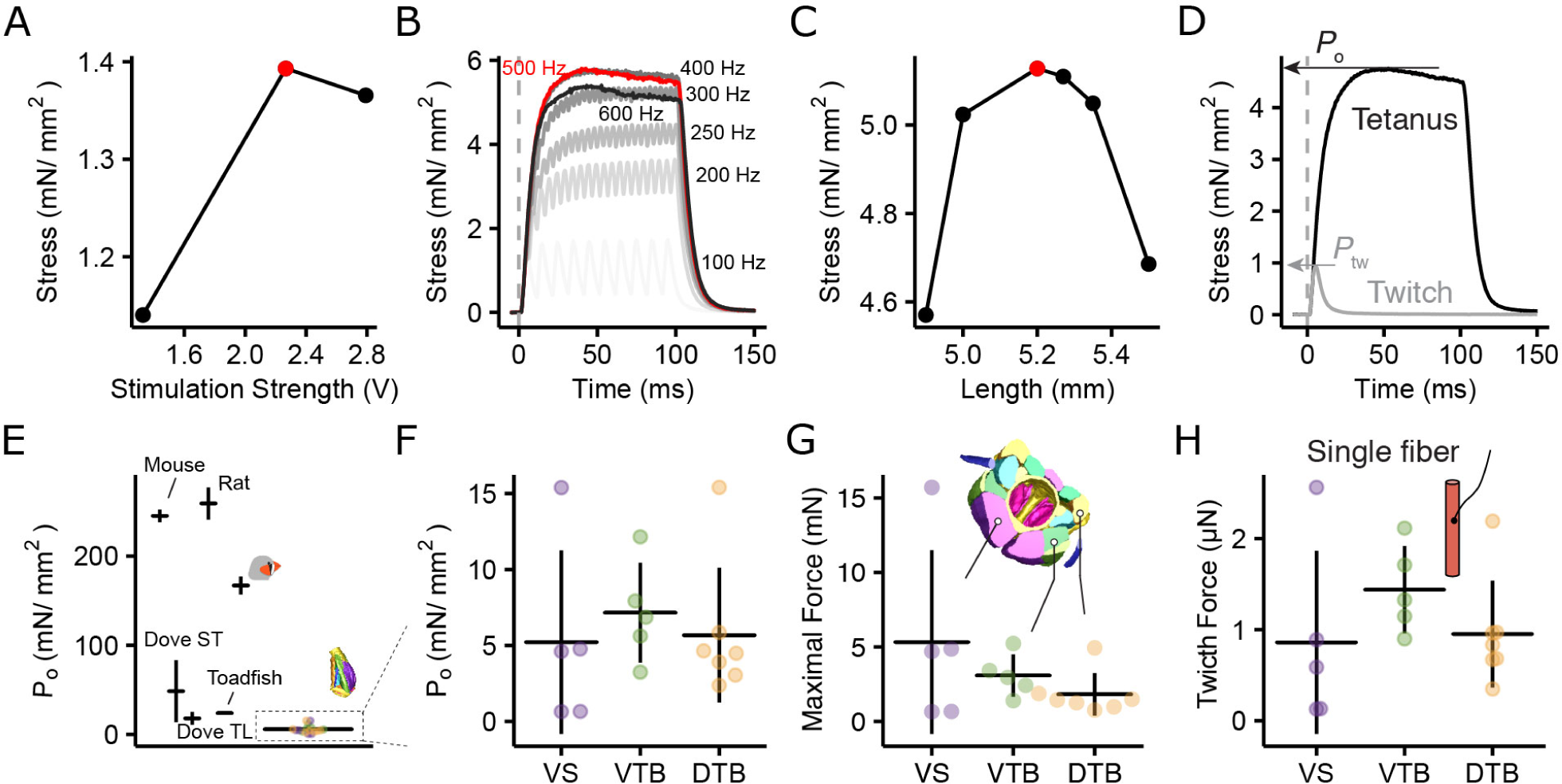
Songbird syringeal muscles generate the lowest stress. Optimization of A) stimulus amplitude, B) frequency and C) muscle fiber length to determine *P*_*o*_. D) Force profile after single (twitch) and high frequency (500 Hz) stimulation. Traces are the mean of eight twitch and three 500 Hz stimulations. E) *P*_*o*_ for syringeal muscles is 30-60x lower than skeletal muscles. VTB; *musculus tracheobonchealis ventralis*, DTB; *musculus tracheobonchealis dorsalis*. F) *P*_*o*_ does not differ significantly between muscles (ANOVA: F = 0.24, df = 2 and 14, p = 0.79) G) Maximal force per muscle. H) Single fiber twitch force is not significantly different between muscles (ANOVA: F = 0.997, df = 2 and 14, p = 0.39)

We measured stress generated by syringeal muscles on isolated fiber bundles *in vitro* for twitch (*P*_tw_) and tetanic (*P*_o_) contractions after a series of length and stimulus amplitude and frequency optimizing experiments (Fig 2A-D). We focused on the ventral syringeal muscle (VS) that predominantly controls fundamental frequency (*f*_o_) or pitch [32-34]. Because VS fibers proved difficult to isolate, we added two other syringeal muscles that control airflow though the syrinx [35]. *P*_o_ did not differ between syringeal muscles and measured 5.21 ± 6.04 mN/mm^2^ (range = 0.64 – 15.4 mN, n = 5) for VS and 5.97 ± 4.46 mN/mm^2^ (range = 0.64 – 15.4 mN, n = 17) for all muscles combined (Fig 2F). Combining *P*_tw_, *P*_o_, cross-sectional area (CSA) and the number of muscle fibers for each muscle (Table S1) provides the maximum force (tetanic stimulation) produced by the entire muscle (Fig 2G) and the minimum force (single twitch) per single muscle fiber (Fig 2H). As such, on average VS force ranged from an average of 0.86 ± 1.00 µN (single twitch in a single fiber) up to 5.31 ± 6.16 mN (full tetanic contraction of the entire muscle). The *P*_o_ of syringeal muscles is 2 and 3 times lower than *P*_o_ of superfast muscles found in bat (9.4 ± 4.2 mN/mm^2^) (See Methods) and toadfish (15 – 24 mN/mm^2^)[36, 37], 4 times lower than pigeon syrinx muscles [29], 30 times lower than zebra finch flight muscles [38], and up to 60 times lower than mammalian limb muscles [30, 31]. Thereby syrinx muscles have the lowest *P*_o_ of any vertebrate skeletal muscle to our knowledge.

### Syringeal MUs provide sub-Hertz resolution control of fundamental frequency

To study the effect of MU recruitment on vocal output, we focused on the control of *f*_o_, an important cue in vocal communication [11]. In birds, analogous to mammalian vocal fold vibrations, expiratory airflow from the bronchi induces self-sustained vibration of vocal fold-like structures, the labia, within the syrinx [39]. In zebra finches, radiated sound pulses are tightly associated with labial collision within the vibration cycle and labial vibration frequency thus directly sets *f*_o_ [33]. When VS shortens, it lengthens the labia and nonlinearly increases their stiffness perpendicular to the expiratory airflow [12]. Together, this stiffness and length increase change the labial resonance frequency, which predominantly increases the labial vibration frequency [10, 12, 40]. By these mechanisms VS force controls *f*_o_, and indeed VS multiunit EMG activity *in vivo* positively correlates to *f*_o_ [32, 34], and VS stimulation *ex vivo* causes *f*_o_ increase [33]. However, it is unknown how the MU force distribution in VS drives changes in *f*_o_.

To quantify the effect of VS force on *f*_o_, we developed an *in vitro* paradigm that allowed for servo-controlled actuation of muscle insertion sites, combined with simultaneous measurement of muscle shortening and force, during sound production in the syrinx (Fig 3A, See Methods). Labial vibration and sound production were induced by increased bronchial and air sac pressure, while actuating the VS insertion site up to 12% shortening of the VS muscle length (Fig 3B) [12]. Frequency transforms linearly with VS force over the full VS force range of 0 – 5.31 mN in three out of five preparations (Fig 3C, Table S2, See methods), and over 65 – 86% of the full VS force range of the remaining two. The linear slopes range from 5 – 57 Hz/mN (mean: 26.6 ± 19.4 Hz/mN, n = 5, Table S2). Because the mechanical properties of the labia are nonlinear [12, 41], this linearization of force to *f*_o_ transformation is surprising and perhaps due to boundary conditions at the edges. We propose this linearization as a beautiful example of morphological computation, where the body locally computes solutions to simplify motor control [42].

**Figure 3.**
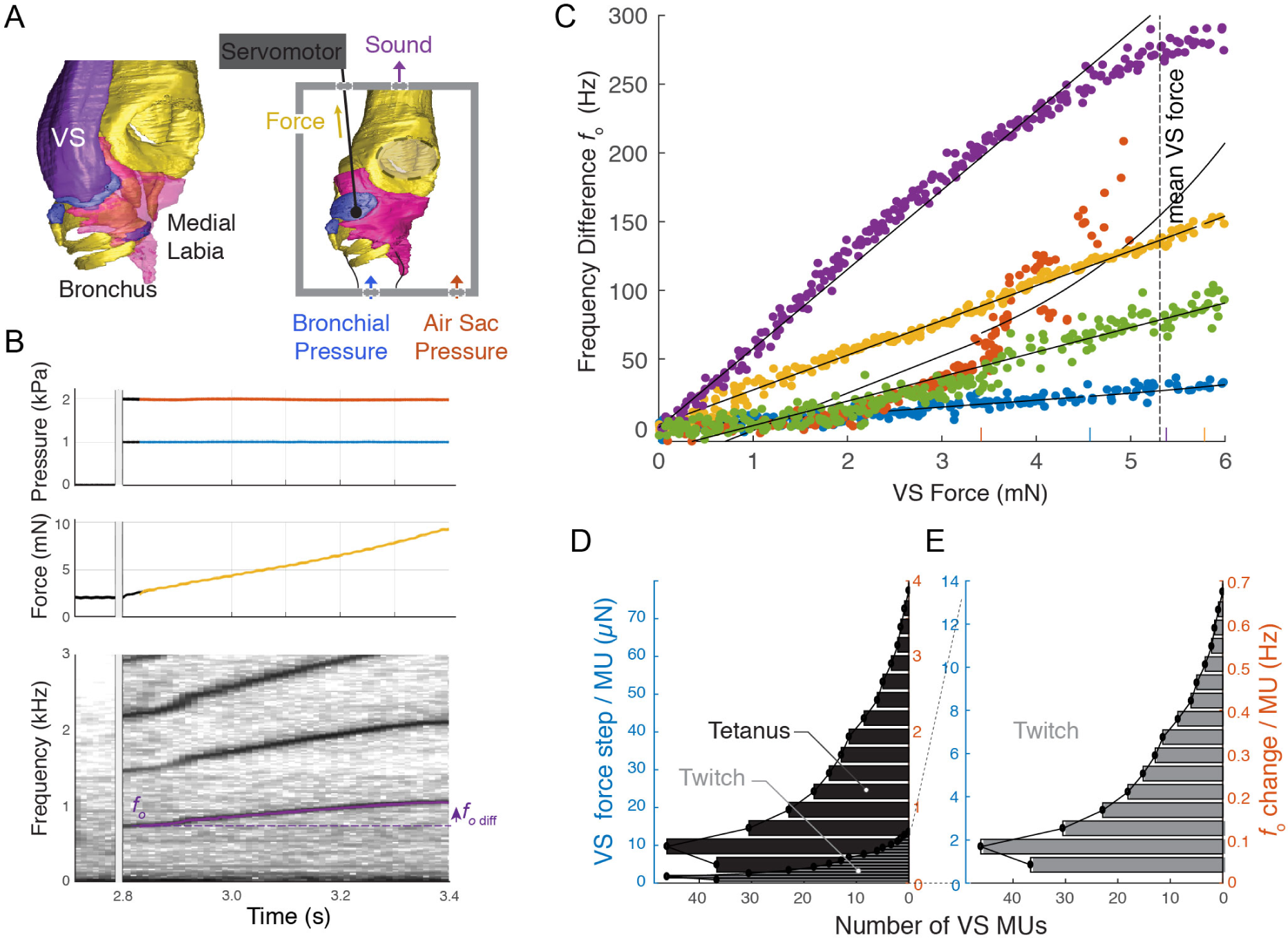
Syringeal MUs provide sub-Hertz resolution control of fundamental frequency. A) Sound production paradigm *in vitro* actuating the insertion site of the *f*_o_ controlling ventral syringeal (VS) muscle. B) Example raw data of sound production induced by raising pressure (top), slow VS force modulation (middle) and resulting *f*_o_ changes (bottom). C) Frequency change as a function of VS force (n = 5) is linear over the VS force modulation range of 0 – 5.31 mN (See Table S2). Short vertical lines indicate statistical breakpoints between linear and exponential curve fits. Colors identify individual birds. D) Distribution of the number of MUs within VS per force and *f*_o_ step available during maximal (tetanic) and E) minimal (twitch) force development.

Combining the MU distribution, *P*_tw_, and *P*_o_ with the muscle force to *f*_o_ transform, we estimated the force and *f*_o_ distribution available to zebra finches (Fig 3D, E). Recruitment of the largest, single, MU containing 15 muscle fibers by full tetanic contraction leads to a mean VS force and *f*_o_ increase of 73 µN and 4.2 Hz, respectively (Fig 3D). Recruiting one of 37 MUs containing a single muscle fiber with a single spike leads to a mean VS force increase of a mere 0.85 µN and *f*_o_ increase of 50 mHz. Maximally recruiting all smallest MUs provides the finest *f*_o_ gradation within a range of 10 Hz, and all second smallest MUs a range of 26 Hz. Taken together, we show that the syringeal muscle motor pool has the highest control resolution possible in the vertebrate nervous system. Combined with the lowest muscle specific stress known in any vertebrate muscle, it allows *f*_o_ control from as small as 50 mHz to 4.2 Hz steps during sound production.

### Behavioral selection on resolution drove small MUs

The fine resolution motor control of *f*_o_ our data suggests is behaviorally relevant. Zebra finch [43] and Bengalese finch males [16] can drive *f*_o_ changes below 1 Hz, respectively, by altered auditory feedback, which corresponds to our estimate of full activation of a single muscle fiber MU. Such small *f*_o_ changes could thus be driven by recruitment of one additional single fiber MU. Moreover, zebra finch females are not only capable of detecting acoustic fine structure [44] and small *f*_o_ changes below 1 Hz [45], they importantly also base their mating decision on those [46]. From a motor control perspective, this suggests a strong selection for small force steps, which can be achieved by small MU sizes combined with low specific force. As a result of the selection on muscle speed, muscle specific force reduces significantly due to architectural constraints of skeletal muscles [27]. Thus, if the force per MU should remain constant and specific force reduced 20 times, MUs could have become 20 times larger to compensate for the force-speed trade-off. However, instead we observe extremely small MUs (13 – 17% of all MUs are innervated one-to-one), which strongly suggests an additional selection for small MUs.

MU innervation numbers below 10 are on the extremely low end of MU sizes in vertebrates, and have so far only been reported for laryngeal [19, 47], extraocular [20, 24] and ear muscles [25]. Interestingly, all those muscles belong to the craniofacial lineage [27] suggesting that their developmental origin might predispose them or even provide unique access to low MU sizes. Thus, developmental origin may in part explain the ability to achieve fine force gradation through small MUs.

### Fine-resolution motor control in songbirds allows vocal space differentiation

Songbirds have radiated explosively ∼40 MYA ago [48], which has been attributed to two key events. First, the evolution of syrinx morphology that may have allowed uncoupled control of specific acoustic features, such as *f*_o_ and amplitude [9, 49], thereby vastly increasing the possibilities or feature space of syllables produced. Second, the evolution of specialized neural circuitry that allowed vocal imitation by trial-and-error learning [3] providing means to explore the vast control space. However, to precisely control their vocal organ to execute trial-to-trial variability, songbirds need fine gradation of force. The songbird syrinx morphology is highly conserved [50] and has comparable numbers of muscle fibers [17] and motor neurons [51] across a wide range of songbird species, which suggests that all songbirds have access to small MUs. We suggest that small MUs were a crucial third key innovation to allow for the fine control of song, one that was pivotal to successfully expanding the acoustic feature space of songbirds, and impetus for the adaptive radiation of today’s roughly 5,000 species of songbirds.

## Acknowledgments

The authors wish to thank Torben Christensen, Per Martensen, Sonja Jacobsen and Bianca Jørgensen for technical support, and Samuel Sober for comments on the manuscript.

## Author Contributions

Conceptualization, IA, CPHE; Methodology, IA, AM, CPHE; Software, IA, CPHE; Formal Analysis, IA, AM, CPHE; Investigation, IA, AM, HR, EBH, MV, CPHE; Writing – Original Draft, CPHE; Writing – Review & Editing, IA, AM, CPHE; Funding Acquisition, IA, CPHE; Resources, IA, CPHE; Supervision, CPHE.

## Material and Methods

### Animal use and care

Adult male zebra finches (*Taeniopygia guttata)* were kept in group aviaries at the University of Southern Denmark, Odense, Denmark on a 12 h light:dark photoperiod and given water and food *ad libitum*. All experiments were conducted in accordance with the Danish law concerning animal experiments and protocols were approved by the Danish Animal Experiments Inspectorate (Copenhagen, Denmark).

### Syrinx extraction

Animals were euthanized by Isoflurane overdose (Baxter, Lillerød, Denmark). The syrinx was dissected out through a ventral incision along the sternum, with regular flushing with oxygenated Ringer’s solution, and submerged in a bath of oxygenated Ringer’s on ice upon removal [29].

### Muscle fiber and motor unit counts

To measure the innervation ratio of syrinx muscles, we counted the number of muscle fibers and axons in the syrinx and tracheosyringeal branch of the hypoglossal nerve (NXIIts) of 8 adult male zebra finches. NXIIts also contains a fraction of sensory fibers, but as they only make up 1% of all axons, we did not correct for them [18].

The syrinx was fixed in 4% PFA in PBS (w/v) for 24 h while keeping the rostro-caudal axis straight. Subsequently it was kept in PBS for 24 h and then embedded in Tissue-Tek O.C.T compound (Sakura Finetek), frozen and stored at −80°C until further processing. All specimen were cut into 10 µm serial cross-sections using a cryotome (Leica CM1860). Immunostainings were performed according to standard protocols using primary antibodies raised against Laminin (10 µg/ml, polyclonal rabbit-anti-Laminin, Sigma-Aldrich Cat# L9393, RRID: AB_477163) and Neurofilament (5 µg/ml, monoclonal mouse-anti-Neurofilament, Millipore Cat# CBL212, RRID: AB_93408) to delineate fiber-boundaries and axons, respectively. Co-labeling was visualized using donkey-anti-mouse-Alexa-Fluor-568 (Abcam Cat# ab175700) and donkey-anti-rabbit-Alexa-Fluor-488 (Abcam Cat# ab150061, RRID: AB_2571722). Slides were coverslipped with Vectasshield mounting medium (Vector Laboratories Cat# H-1000, RRID: AB_2336789) and sealed with transparent nail polish. Images were acquired with a Zyla sCMOS camera (AndorTM Technology Ltd, Northern Ireland) mounted on a Nikon Eclipse T*i* inverted microscope with automated XY-stage. To count the number of muscle fibers, we acquired images from sections with a 20x objective directly rostral of bronchial bone B2, a cross-sectional plane where all muscle fibers are present. To cover the entire syrinx cross-section, we acquired 600-800 images in random order and stitched them together with NIS-Elements (Nikon). Muscles were counted on 10x bicubic down-sampled images (minimal image size: 5314×4293 pixels). Axons were counted in images acquired with a 100x objective from sections in the region of tracheal ring T4-T6 (minimal image size: 2561×3996 pixels). Counts were presented as the averaged of counts by three independent observers (IA, AM, BJ) using the ImageJ (ImageJ, RRID:SCR_003070) Cell Counter plugin.

### MU size distribution estimates

We exploited the fact that the frequency distribution of MU size or MU force is consistently skewed for all muscles studied to date [21-23] to estimate the distribution of MUs within syringeal muscles. The MU size frequency distribution follows the exponential curve:

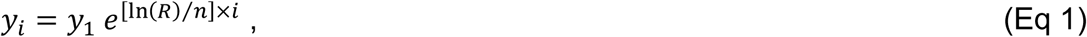

where *y*_i_ is the innervation number of MU *i, y*_1_ is the innervation number for the smallest MU (unit 1), *n* is the number of MUs, and *R* is the ratio of innervation numbers for the largest and smallest unit: *R* = *y*_*n*_/*y*_1_ [52]. This relation allowed us to calculate the number of muscle fibers within the muscle (*N*_MF_) for a given maximum MU size as 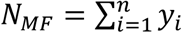. Thus, by knowing *N*_MF_ for a muscle, we can calculate the maximum MU size and MU frequency distribution.

### Muscle specific tension

We characterized muscle specific tension of the *musculus syringealis ventralis* (VS; n = 5), *musculus tracheobonchealis ventralis* (VTB; n =5), and *musculus tracheobonchealis dorsalis* (DTB; n = 7) of adult male zebra fiches on preparations of isolated muscle fiber bundles as previously described [53]. In brief, fiber bundles were mounted in a temperature-controlled bath, which was continuously supplied with oxygenated Ringers solution. The rostral end of the preparation was fixed to a force transducer (Model 400A, Aurora Scientific) and the caudal end to a micromanipulator, which was used to control length of the preparation. Field stimulations were applied using platinum electrodes. Force and stimulation signals were low pass filtered (EF120 BNC through-feed low-pass filter, Thor Labs) and digitized at 40 kHz (NI DAQ Board PCI-MIO-16E4, National Instruments). The force baseline was defined as the average amplitude of 50 ms of the force signal prior to stimulation onset and subtracted from all force data.

To measure *P*_o_, we optimized first *s*timulation amplitude (at pulse width of 300 µs) for maximal force production, followed by tetanic force frequency curve by 100 ms pulse trains ranging from 100 to 800 Hz in 100 Hz steps, and finally resting length L_o_. Isometric stress was calculated as F/A_csa_ of the muscle, where the cross-sectional area A_csa_ was estimated from the resting length L_o_ and the dry weight (dry-wet conversion factor: 5) of the muscle fibers assuming a constant density of 1060 kg/m^3^ from [54]. Muscle specific force was calculated using the average CSA as in Table S1. Similarly, the force per muscle fiber was obtained by dividing the muscle specific force by the average number of muscle fibers as in Table S1.

We determined the *P*_o_ of bat superfast laryngeal muscle by re-analyzing previously published data [55].

All control software was written in MATLAB (MathWorks, RRID:SCR_001622) and all analyses were performed in R (R Project for Statistical Computing, RRID:SCR_001905).

### In vitro sound production

We developed an experimental paradigm that allowed for sound production *in vitro* building on earlier work. Previously, we induced syringeal sound production by precisely controlling bronchial and air sac pressure *in vitro* or in perfused preparations *ex vivo* [33]. Additionally, we developed methods to actuate muscle insertion sites but without inducing sound [12]. Here we further developed our experimental chamber and combined sound induction by pressure control with the actuation of a single syringeal muscle insertion site and force measurements. In brief, directly after syrinx extraction, the *musculus syringealis ventralis* (VS) was removed while the syrinx was still submerged in Ringers solution. The syrinx was then moved to the experimental chamber and a 10–0 suture needle was inserted through the center of the bottom edge of the *medio-ventral cartilage* (MVC) pads. The mono-filament suture was threaded through a 100 µm hole and attached to a servomotor (Ergometer model 322C, Aurora Scientific) that controlled length while measuring force at the tip of the lever arm (displacement and force resolution 1 μm and 0.3 mN, respectively). To induce sound, we applied pressure differential over the syringeal labia (air sac pressure 2.0 kPa, bronchial pressure 1.0 kPa) using dual-valve differential pressure PID controllers (model PCD, 0–10 kPa, Alicat Scientific). Next, we actuated the MVC in a 500 ms ramp from 0 to maximally 12% of the VS length [12], and measured the corresponding shift in *f*_o_ in 5 syrinx preparations.

Sound was recorded with a ½-inch pressure microphone-pre-amplifier assembly (model 46AD with preamplifier type 26AH, G.R.A.S., Denmark), amplified and high-pass filtered (10 Hz, 3-pole Butterworth filter, model 12AQ, G.R.A.S.). The microphone sensitivity was measured before each experiment (sound calibrator model 42AB, G.R.A.S.). The microphone was placed at 2 – 3 cm from the tracheal connector outlet in the acoustic near field, and on a 45° angle to avoid the air jet from the tracheal outlet.

Microphone, pressure, force, and displacement signals were low-pass filtered at 10 kHz (EF120 BNC through-feed low-pass filter, Thor Labs) and digitized at 50 kHz (USB 6259, 16 bit, National Instruments). We calculated the mean values for pressure, force and displacement signals in 2 ms bins with a 1 ms sliding window. To calculate *f*_o_ for each bin, we used the Yin algorithm [56] within a frequency range of 350 – 1300 Hz, and aperiodicity threshold between 0.1 – 0.2. All control and analysis software was written in Matlab (MathWorks).

### Statistics

All analyses were performed in MATLAB (MathWorks, RRID:SCR_001622) or R (R Project for Statistical Computing, RRID:SCR_001905). All values are presented as mean ± SD. The Welch Two Sample t-test was used to test for significant differences between left and right NXIIts axon and muscle fiber numbers. Root innervation numbers were pooled across left and right syrinx halves when not significantly different (p>0.05). Statistical difference between force production by VS, VTB and DTB muscles was assessed using ANOVA.

The VS force (*x*) -*f*_o_ (*y*) transformation was modeled with a linear model at low force and an exponential model at high force [57]:

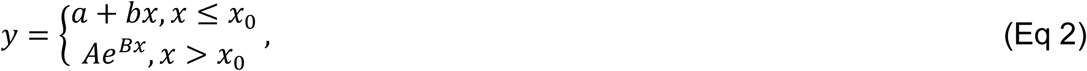

where *x*_0_ is the linear limit. To satisfy the continuous and differentiable requirements, we set 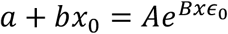 and 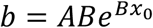, and thus the linear limit is 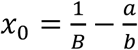. The parameters were determined by iteratively optimizing the model fit minimizing least squares using the MATLAB routine *lsqcurvefit*.

## Supplemental Information titles and legends

**Table S1.**
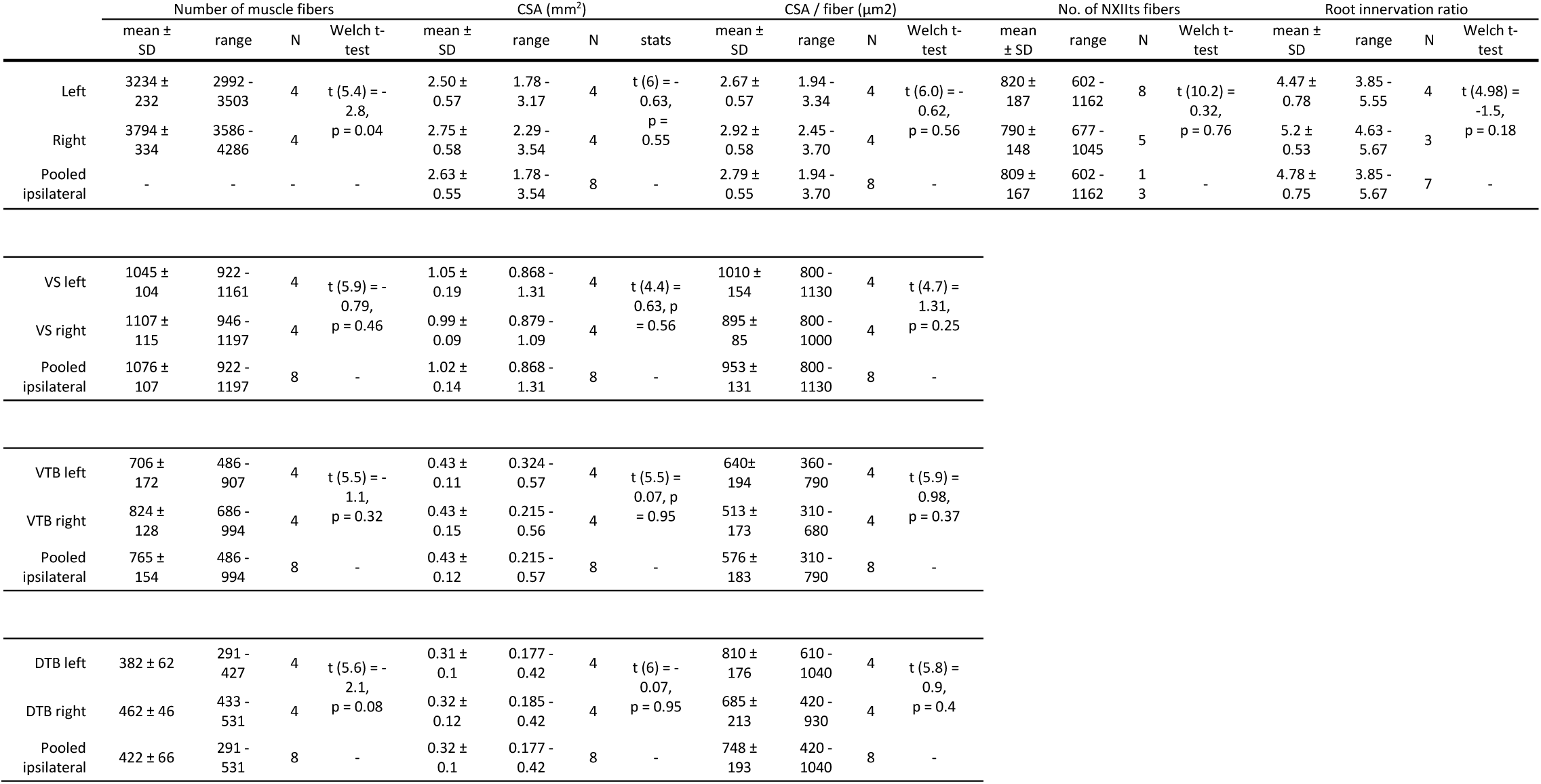
Syringeal muscle and NXIIts nerve properties in the adult male zebra finch.

**Table S2.** Linear and exponential model fit parameters of force-*f*_o_ transformation. The fitted function was of the form: 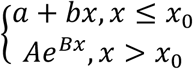.

| Individual | Linear Fit<br>y=ax+b |  | Exponential Fit<br>y=Ae<sup>(Bx)</sup> |  | x<sub>0</sub> |  |
| --- | --- | --- | --- | --- | --- | --- |
|  | a | b | A | B | (mN) | % of VS max |
| 1 | 5.0 | 0.0 | 8.4 | 0.22 | 4.57 | 86 |
| 2 | 27.1 | -29.0 | 16.0 | 0.42 | 3.42 | 64 |
| 3 | 25.3 | 2.2 | 55.4 | 0.17 | 5.78 | >100 |
| 4 | 57.6 | 0.1 | 114.0 | 0.19 | 5.38 | >100 |
| 5 | 17.8 | -16.3 | 46.4 | 0.13 | 8.80 | >100 |
| mean + SD | 26.6 ± 19.4 | -8.6 ± 13.6 | 48.0 ± 41.8 | 0.23 ± 0.12 | 5.6 ± 2.0 |  |

## References

1. Enoka, R.M., and Fuglevand, A.J. (2001). Motor unit physiology: some unresolved issues. Muscle Nerve 24, 4–17.

2. Raikova, R., Celichowski, J., Angelova, S., and Krutki, P. (2018). A model of the rat medial gastrocnemius muscle based on inputs to motoneurons and on an algorithm for prediction of the motor unit force. J Neurophysiol 120, 1973–1987.

3. Fee, M.S., and Scharff, C. (2010). The songbird as a model for the generation and learning of complex sequential behaviors. Ilar J 51, 362–377.

4. Vicario, D.S., and Nottebohm, F. (1988). Organization of the zebra finch song control system: I. Representation of syringeal muscles in the hypoglossal nucleus. J Comp Neurol 271, 346–354.

5. Tang, C., Chehayeb, D., Srivastava, K., Nemenman, I., and Sober, S.J. (2014). Millisecond-scale motor encoding in a cortical vocal area. PLoS Biol 12, e1002018.

6. Price, T. (1998). Sexual selection and natural selection in bird speciation. Philos T R Soc B 353, 251–260.

7. Uy, J.A.C., Irwin, D.E., and Webster, M.S. (2018). Behavioral Isolation and Incipient Speciation in Birds. Annu Rev Ecol Evol S 49, 1–24.

8. Janik, V.M., and Slater, P.J. (2000). The different roles of social learning in vocal communication. Anim Behav 60, 1–11.

9. Gaunt, A.S. (1983). An Hypothesis Concerning the Relationship of Syringeal Structure to Vocal Abilities. Auk 100, 853–862.

10. Titze, I., Riede, T., and Mau, T. (2016). Predicting Achievable Fundamental Frequency Ranges in Vocalization Across Species. PLoS Comput Biol 12, e1004907.

11. Goller, F., and Riede, T. (2013). Integrative physiology of fundamental frequency control in birds. J Physiol Paris 107, 230–242.

12. During, D.N., Knorlein, B.J., and Elemans, C.P.H. (2017). In situ vocal fold properties and pitch prediction by dynamic actuation of the songbird syrinx. Sci Rep 7, 11296.

13. Adam, I., and Elemans, C.P.H. (2019). Vocal Motor Performance in Birdsong Requires Brain-Body Interaction. eNeuro 6.

14. Xiao, L., Chattree, G., Oscos, F.G., Cao, M., Wanat, M.J., and Roberts, T.F. (2018). A Basal Ganglia Circuit Sufficient to Guide Birdsong Learning. Neuron 98, 208–221 e205.

15. Tumer, E.C., and Brainard, M.S. (2007). Performance variability enables adaptive plasticity of ‘crystallized’ adult birdsong. Nature 450, 1240–1244.

16. Sober, S.J., and Brainard, M.S. (2009). Adult birdsong is actively maintained by error correction. Nat Neurosci 12, 927–931.

17. Christensen, L.A., Allred, L.M., Goller, F., and Meyers, R.A. (2017). Is sexual dimorphism in singing behavior related to syringeal muscle composition? The Auk 134, 710–720.

18. Lissandrello, C.A., Gillis, W.F., Shen, J., Pearre, B.W., Vitale, F., Pasquali, M., Holinski, B.J., Chew, D.J., White, A.E., and Gardner, T.J. (2017). A micro-scale printable nanoclip for electrical stimulation and recording in small nerves. J Neural Eng 14, 036006.

19. Santo, H., and Marques, M.J. (2008). Estimation of the number and size of motor units in intrinsic laryngeal muscles using morphometric methods. Clin Anat 21, 301–306.

20. Mühlendyck, H. (1978). The Size of Motor Units in Reference to Eye-Muscle Fibres of Different Innervation. (Munich: J.F. Bergmann-Verlag), pp. 17–26.

21. Olson, C.B., and Swett, C.P. (1966). A Functional and Histochemical Characterization of Motor Units in a Heterogeneous Muscle (Flexor Digitorum Longus) of Cat. Journal of Comparative Neurology 128, 475-&.

22. Milner-Brown, H.S., Stein, R.B., and Yemm, R. (1973). The orderly recruitment of human motor units during voluntary isometric contractions. J Physiol 230, 359–370.

23. Elek, J.M., Kossev, A., Dengler, R., Schubert, M., Wohlfahrt, K., and Wolf, W. (1992). Parameters of Human Motor Unit Twitches Obtained by Intramuscular Microstimulation. Neuromuscular Disord 2, 261–267.

24. Bruenech, J.R., and Kjellevold Haugen, I.B. (2015). How does the structure of extraocular muscles and their nerves affect their function? Eye (Lond) 29, 177–183.

25. Lu, J., Tapia, J.C., White, O.L., and Lichtman, J.W. (2009). The interscutularis muscle connectome. PLoS Biol 7, e32.

26. Elemans, C.P., Mead, A.F., Rome, L.C., and Goller, F. (2008). Superfast vocal muscles control song production in songbirds. PLoS One 3, e2581.

27. Mead, A.F., Osinalde, N., Ortenblad, N., Nielsen, J., Brewer, J., Vellema, M., Adam, I., Scharff, C., Song, Y., Frandsen, U., et al. (2017). Fundamental constraints in synchronous muscle limit superfast motor control in vertebrates. Elife 6.

28. Rome, L.C., and Lindstedt, S.L. (1998). The Quest for Speed: Muscles Built for High-Frequency Contractions. News Physiol Sci 13, 261–268.

29. Elemans, C.P., Spierts, I.L., Hendriks, M., Schipper, H., Muller, U.K., and van Leeuwen, J.L. (2006). Syringeal muscles fit the trill in ring doves (Streptopelia risoria L.). J Exp Biol 209, 965–977.

30. Brooks, S.V., and Faulkner, J.A. (1991). Forces and powers of slow and fast skeletal muscles in mice during repeated contractions. J Physiol 436, 701–710.

31. Urbanchek, M.G., Picken, E.B., Kalliainen, L.K., and Kuzon, W.M., Jr. (2001). Specific force deficit in skeletal muscles of old rats is partially explained by the existence of denervated muscle fibers. J Gerontol A Biol Sci Med Sci 56, B191–197.

32. Srivastava, K.H., Elemans, C.P., and Sober, S.J. (2015). Multifunctional and Context-Dependent Control of Vocal Acoustics by Individual Muscles. J Neurosci 35, 14183–14194.

33. Elemans, C.P., Rasmussen, J.H., Herbst, C.T., During, D.N., Zollinger, S.A., Brumm, H., Srivastava, K., Svane, N., Ding, M., Larsen, O.N., et al. (2015). Universal mechanisms of sound production and control in birds and mammals. Nat Commun 6, 8978.

34. Goller, F., and Suthers, R.A. (1996). Role of syringeal muscles in controlling the phonology of bird song. J Neurophysiol 76, 287–300.

35. Goller, F., and Suthers, R.A. (1996). Role of syringeal muscles in gating airflow and sound production in singing brown thrashers. J Neurophysiol 75, 867–876.

36. Rome, L.C., Cook, C., Syme, D.A., Connaughton, M.A., Ashley-Ross, M., Klimov, A., Tikunov, B., and Goldman, Y.E. (1999). Trading force for speed: why superfast crossbridge kinetics leads to superlow forces. Proc Natl Acad Sci U S A 96, 5826–5831.

37. Young, I.S., and Rome, L.C. (2001). Mutually exclusive muscle designs: the power output of the locomotory and sonic muscles of the oyster toadfish (Opsanus tau). Proceedings. Biological sciences / The Royal Society 268, 1965–1970.

38. Ellerby, D.J., and Askew, G.N. (2007). Modulation of flight muscle power output in budgerigars Melopsittacus undulatus and zebra finches Taeniopygia guttata: in vitro muscle performance. J Exp Biol 210, 3780–3788.

39. Düring, D.N., and Elemans, C.P.H. (2016). Embodied Motor Control of Avian Vocal Production. In Vertebrate Sound Production and Acoustic Communication, R.A. Suthers, W.T. Fitch, R.R. Fay and A.N. Popper, eds. (Cham: Springer International Publishing), pp. 119–157.

40. Riede, T., and Goller, F. (2010). Functional morphology of the sound-generating labia in the syrinx of two songbird species. J Anat 216, 23–36.

41. Fee, M.S. (2002). Measurement of the linear and nonlinear mechanical properties of the oscine syrinx: implications for function. J Comp Physiol A Neuroethol Sens Neural Behav Physiol 188, 829–839.

42. Pfeifer, R., Lungarella, M., and Iida, F. (2007). Self-organization, embodiment, and biologically inspired robotics. Science 318, 1088–1093.

43. Andalman, A.S., and Fee, M.S. (2009). A basal ganglia-forebrain circuit in the songbird biases motor output to avoid vocal errors. Proc Natl Acad Sci U S A 106, 12518–12523.

44. Prior, N.H., Smith, E., Lawson, S., Ball, G.F., and Dooling, R.J. (2018). Acoustic fine structure may encode biologically relevant information for zebra finches. Sci Rep-Uk 8.

45. Lohr, B., and Dooling, R.J. (1998). Detection of changes in timbre and harmonicity in complex sounds by zebra finches (Taeniopygia guttata) and budgerigars (Melopsittacus undulatus). J Comp Psychol 112, 36–47.

46. Woolley, S.C., and Doupe, A.J. (2008). Social context-induced song variation affects female behavior and gene expression. PLoS Biol 6, e62.

47. Hinrichsen, C.F., and Ryan, A. (1982). The size of motor units in laryngeal muscles of the rat. Experientia 38, 360–361.

48. Feduccia, A. (2003). ‘Big bang’ for tertiary birds? Trends in Ecology & Evolution 18, 172–176.

49. During, D.N., Ziegler, A., Thompson, C.K., Ziegler, A., Faber, C., Muller, J., Scharff, C., and Elemans, C.P. (2013). The songbird syrinx morphome: a three-dimensional, high-resolution, interactive morphological map of the zebra finch vocal organ. BMC Biol 11, 1.

50. King, A. (1989). Functional anatomy of the syrinx. In Form and Function in Birds, Volume 4, A. King and J. McLelland, eds. (London, UK: Academic Press), pp. 105–192.

51. Moore, J.M., Szekely, T., Buki, J., and Devoogd, T.J. (2011). Motor pathway convergence predicts syllable repertoire size in oscine birds. Proc Natl Acad Sci U S A 108, 16440–16445.

52. Fuglevand, A.J., Winter, D.A., and Patla, A.E. (1993). Models of recruitment and rate coding organization in motor-unit pools. J Neurophysiol 70, 2470–2488.

53. Srivastava, K.H., Holmes, C.M., Vellema, M., Pack, A.R., Elemans, C.P.H., Nemenman, I., and Sober, S.J. (2017). Motor control by precisely timed spike patterns. P Natl Acad Sci USA 114, 1171–1176.

54. Mendez, J., and Keys, A. (1960). Density and Composition of Mammalian Muscle. Metabolism 9, 184–188.

55. Elemans, C.P., Mead, A.F., Jakobsen, L., and Ratcliffe, J.M. (2011). Superfast muscles set maximum call rate in echolocating bats. Science 333, 1885–1888.

56. de Cheveigne, A., and Kawahara, H. (2002). YIN, a fundamental frequency estimator for speech and music. J Acoust Soc Am 111, 1917–1930.

57. Zhang, Y.S., Takahashi, D.Y., Liao, D.A., Ghazanfar, A.A., and Elemans, C.P.H. (2019). Vocal state change through laryngeal development. Nat Commun 10, 4592.

